# Transcriptomics and mutational analysis to screen immunogenic neoantigen peptides and Patient stratification based on immune subtypes for TNBC

**DOI:** 10.64898/2026.02.18.706559

**Authors:** Karthick Vasudevan, T Dhanushkumar, Prasanna Kumar Selvam, Anbarasu Krishnan, B G Sunila, Santhosh Mudipalli Elavarasu, Supraja Mohan, Rohini Karunakaran

## Abstract

Triple-negative breast cancer (TNBC) is a highly aggressive and heterogeneous subtype with limited therapeutic options. In this study, we performed an integrative analysis of TNBC genomics data, including gene expression, somatic mutations, copy number alterations, survival outcomes, immune profiling, and clustering, to identify potential neoantigens, patient populations suitable for vaccination, and biomarkers for evaluating vaccine efficacy. This Integrated analysis identified POSTN and CAP1 as tumor-specific antigens. Incorporation of TNBC-specific mutations into the screened wild-type antigens led to the identification of three neoantigenic peptides with high potential for vaccine development. Immune subtyping stratified TNBC patients into four distinct subtypes, among which IS1 and IS3 were characterized by poor immune infiltration, lower mutation burden, and unfavorable prognosis, whereas IS2 and IS4 exhibited enhanced immune activity and better clinical outcomes. A vaccine incorporating the identified neoantigen peptides may potentially remodel the immune landscape of immune-cold subtypes (IS1 and IS3), converting them into immune-enriched phenotypes through vaccine-induced immune stimulation. Furthermore, weighted gene co-expression network analysis identified ten immune-related biomarkers from the blue and gray modules that were significantly associated with improved survival in IS2 and IS4. Functional enrichment and protein–protein interaction analyses revealed that hub genes primarily involved in immunoglobulin kappa chains and cytokine/TNF signaling pathways may serve as valuable immune biomarkers for prognostic assessment and monitoring vaccine efficacy.

## 1. Introduction

Triple-negative breast Cancer (TNBC) is an aggressive and specific subtype of breast cancer that lacks Estrogen Receptors (ER), Progesterone Receptors (PR), and Human Epidermal Growth Factor Receptor 2 (HER2) [1]. This subtype poses a major issue in therapeutic approaches and shows poor survival rates. TNBC accounts for approximately 10–20% of all breast cancer cases worldwide. This subtype is distinct in its patterns of metastasis and has no targeted therapies yet [2–3]. Compared to other breast cancer subtypes, TNBC cells are highly proliferative and, thus, have a poor prognosis. The lack of hormone receptors severely restricts treatment options. Due to these limitations, surgical removal of the tumour followed by radiotherapy and chemotherapy remained the standard therapy options with their side effects. Researchers are looking for other options to tackle this aggressive subtype [4].

Immunotherapy has reshaped the treatment landscape for cancer, particularly with the rise of immune checkpoint inhibitors (ICIs) like pembrolizumab. While these drugs have become a cornerstone for certain patients, their limitations have sparked innovative strategies to maximize their potential [5]. ICP inhibitors like Programmed Death Ligand 1 inhibitors have very poor survival rates in clinical trials, with only 35% survival rates. This limitation is due to the high heterogeneity observed in TNBC. This limited effectiveness has driven research into combining ICIs with other therapeutic strategies, such as cancer vaccines, to enhance their efficacy and broaden the patient population that benefits from these treatments. [6–8].

A cancer vaccine works by using the body’s immune system capacity to identify and target tumor antigens of tumor cells. Many tumors arise due to the mutation of normal genes. These mutations cause the production of many tumor-specific proteins that are not present in normal cells, which makes them non-self proteins. Immune cells recognize them and neutralize them, however, tumors have adopted many mechanisms for immune evasion, like the downregulation of tumor antigens during immune surveillance, producing immune checkpoints [9–10]. Various forms of therapeutic vaccines are currently being developed, including those based on peptides, carbohydrate-based, genetic, and dendritic cell (DC) vaccines. Each vaccine type has unique mechanisms of action and potential benefits [11–18].

mRNA-based cancer vaccines incorporating immunogenic tumor antigens have emerged as a promising therapeutic strategy due to several distinct advantages. A key benefit lies in their rapid development and scalable manufacturing. This accelerated production cycle is particularly advantageous in oncology, where timely intervention can substantially influence clinical outcomes. Another major advantage of mRNA vaccines is their capacity for personalization. Unlike conventional cancer vaccines that target broadly expressed shared antigens, mRNA platforms can be tailored to encode tumor-specific mutations (neoantigens) unique to an individual patient’s malignancy [19–20]. This strategy enables a highly specific and robust anti-tumor immune response, as the immune system is directed against non-self epitopes derived from tumor-specific genetic alterations. Such precision is critical in cancer settings where tumors frequently undergo clonal evolution and develop escape mechanisms or therapeutic resistance [21–22]. Importantly, mRNA vaccines exhibit strong potential for combination therapeutic regimens. When administered alongside immune checkpoint inhibitors, they may enhance antigen-specific T-cell responses and overcome immunosuppressive features of the tumor microenvironment, thereby improving clinical efficacy. Recent clinical trials have demonstrated encouraging outcomes with mRNA vaccines across multiple malignancies, including melanoma and other solid tumors. The present study aimed to identify novel neoantigens associated with TNBC and to define distinct immune subtypes within the TNBC population to screen patients suitable for a vaccine, followed by biomarker identification to monitor vaccine effect. Such stratification will facilitate rational patient selection for mRNA vaccine-based immunotherapy and may improve therapeutic responsiveness in this highly aggressive breast cancer subtype.

## 2. Methods and materials

### 2.1. Data collection

The data for this study were obtained from the Cancer Genome Atlas of TCGA through the GDC Data Portal, focusing on TNBC and samples of normal breast tissue. A total of 121 TNBC and 112 normal count files were collected using the GDC Data Transfer Tool [23], specifically RNA sequencing data. Clinical information for these 121 patients was also gathered (patient IDs, last follow-up days, death days, and vital status of all 121 patients are tabulated in the supplementary Table 1). The TNBC samples were selected by applying filters for the negative status of ER, PR, and HER2. Additionally, 103 TNBC MAF files [24] were obtained from the GDC portal for the analysis of mutations in TNBC samples. Additionally, amplified Copy number alteration (CNA) TNBC genes were collected from cBioPortal.

### 2.2. Tumor antigen screening

To assess gene expression and genetic variability in TNBC, differential expression analysis was performed using DESeq2 [25] on the RNA-Seq count files. The analysis compared 121 TNBC samples to 112 normal samples. log2 fold exceeding 1 with a significant p-value (< 0.0005) was utilized to identify genes that were significantly upregulated in the TNBC samples, which were then stored for further analysis. Subsequently, the MAF files containing mutation data from 103 TNBC samples were analysed using the maftools R package [26]. The intersection of upregulated, CNA, and mutated genes was then analysed to identify potential tumor antigens.

### 2.3. Prognostic Survival Outcome Evaluation

To evaluate the prognostic relevance of candidate tumor antigen genes, an overall survival analysis [27] was conducted using the survival library from the R package [28]. Patients were divided into high and low-expression groups for each gene according to the median expression level. Kaplan-Meier (KM) survival curves were constructed to compare overall survival between these two groups. HR values [29] were considered to evaluate the recurrence rate.

### 2.4. Analysis of Antigen-Immune Cell Correlations

The TIMER 2.0 [30] web tool was utilized to investigate the relationship between selected tumor antigens and antigen-presenting immune cells, including macrophages, B cells, and dendritic cells. TIMER uses Spearman’s correlation between input genes with the selected immune cells.

### 2.5. Immune Subtype Profiling in TNBC

For immune subtype profiling in TNBC, a list of 1,753 immune-related genes was retrieved from the ImmPort database [31]. This gene list was utilized to filter the gene expression count data for TNBC, retaining only the immune counts from the count file. The filtered gene expression data were employed for immune subtyping analysis. The analysis was performed using the ConsensusClusterPlus R package, which used a PAM algorithm similar to the k-means method, but this method used real data points [32]. To perform consensus clustering, the maximum K-value was set to nine clusters with 500 bootstraps and 80% of the samples selected for each resampling.

### 2.6. Immune-related molecular features of subtypes and Prognostic analysis of immune subtypes

To assess immune activity across different TNBC subtypes, the C5 GO biological process file which contain 7608 pathways related DNA repair, cell cycle regulation, metabolism, gene expression, protein modification, and immune response, were downloaded from the Molecular Signatures Database of GSEA website, from them 650 pathway relates to immune system were retained [33] and annotated to immune subtype (IS) genes using the FGSEA package in R, and enrichment scores for each immune pathway for each sample were generated [34]. From them, 17 pathways highly related to the immune response were considered for the immune survey analysis. The immune landscape score of all subtypes was visualized using a heatmap according to the expression levels of the chosen immune-related genes in each subtype. The ESTIMATE package of the R [35] was utilized to compute the tumor stromal score, immune score, and estimate scores, offering insights into immune infiltration and stromal content. Visualization of the ESTIMATE results was done using box plots. The CIBERSORT R package [36] was employed to quantify the relative proportions of 22 different immune cell types within the immune subtypes, providing a detailed characterization of the immune cell composition. Following these analyses, overall survival for each subtype was calculated using the survival package in R [37], highlighting the prognostic significance of immune subtypes within TNBC, Immune checkpoint genes (ICP), and distribution of immune cell death genes (ICD) across the different subtype samples. were evaluated and visualized using box plots.

### 2.7. WGCNA to identify the biomarker for mRNA vaccine response

WGCNA analysis was performed to identify hub gene biomarkers responsible for favorable survival rates in patients with high survival rates, using the WGCNA library in R [38]. A soft-thresholding power of 6 was established, and the topological overlap matrix (TOM) was computed to assess the similarity among genes based on their shared connections. Hierarchical clustering was then applied to the TOM-derived dissimilarity matrix, generating a gene tree with a deep split of 3 to detect gene modules through dynamic tree cutting. A minimum threshold of 30 genes was set for each module. Counts for genes in each module from each IS were analysed using a box plot, survival analysis for each module was carried out, and the best HR value module’s biological process, molecular function, and KEGG pathway analysis were carried out using ShinyGO 0.80 [39–40]. The PPI network using the STRING database was constructed for genes of selected modules [41], and using Cytoscape tools, hub genes were calculated for those modules [42].

## 3. Results

### 3.1. Tumor antigen identification

Differential expression analysis revealed 14,776 upregulated genes in the TNBC samples [43–44]. The results are visualized using a heatmap, as illustrated in Figure 1A, which presents the top 50 most differentially expressed genes and a volcano plot showing the differentiation of upregulated, downregulated, and undifferentiated genes, as illustrated in Figure 1B [45]. Mutation analysis identified 6,822 genes the mutated genes were found across a range of samples, with each gene varied in at least 7% of the total sample to 89% The most prevalent mutations were frameshift deletion, Frame Shift Insert, In Frame deletion, In Frame Insert, Missense mutation, and Nonsense mutation, the prevalence and variation of mutations across different genes and samples are shown in Figure 1C, additionally this mutational analysis revealed mutation of TP53 across many samples. Additionally, a total of 18,531 CAN genes were observed in an amplified form. To discover potential tumor antigens, we intersected the upregulated genes, CAN, and mutated genes [46–47]. This intersection yielded 2,264 genes (Figure 1D), considered potential tumor antigens for further analysis. These genes are significantly upregulated with amplified nature in TNBC samples and possess mutations, which suggests that the mutated proteins are over-expressed specifically in TNBC and carry non-self protein, which can be detected by immune cells to induce an immune response [48].

**Figure 1.**
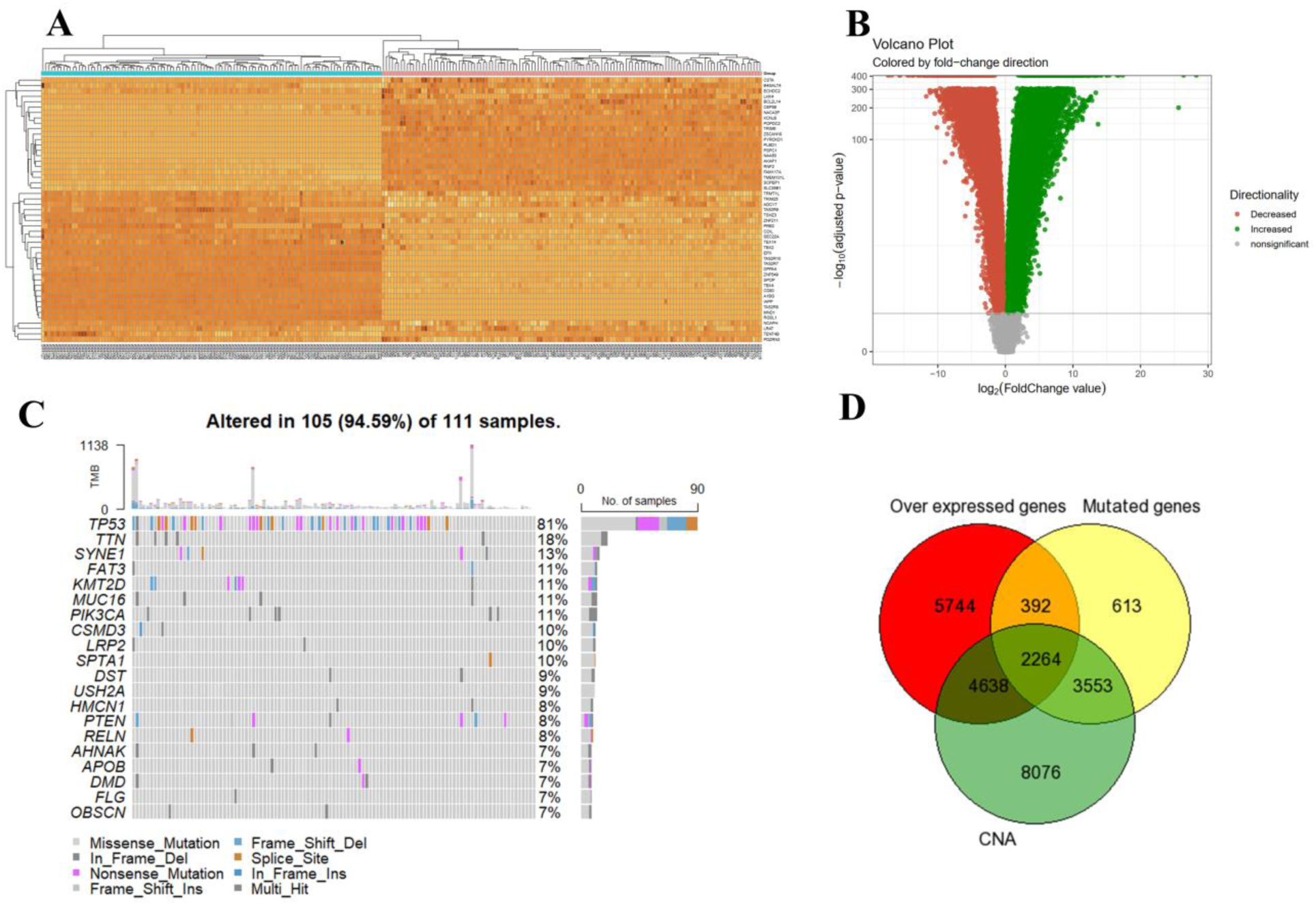
Tumor antigen screening results A) heatmap of top 50 upregulated genes, B) volcano plot, C) Oncoplot representing the mutation in each sample and type of mutation along with the mutation burden in each patient, D) Venn diagram of intersection region of over-expressed, mutated, and CAN genes

### 3.2. Survival Analysis

In comprehensive overall survival analysis of 2,264 tumor antigens revealed divergent p and HR values across these tumor antigens, KM survival analysis revealed seven genes *ALG2, SURF4, CAP1, COL5A3, POSTN, GOLGB1*, and *MFAP5* with statistically significant associations with overall survival, with p-values below 0.005 and HR value less than 1, underscoring their profound impact on patient prognosis [49]. These genes exhibited low HRs, indicating a decreased likelihood of recurrence, suggesting that patients with these genes have better progression [50]. Figure 2A presents a bar plot of the p-values and HRs for the top 50 genes, and Figures 2B to 3H display detailed KM survival plots for the seven most significant genes, illustrating survival differences between patients with elevated and reduced expression levels.

**Figure 2.**
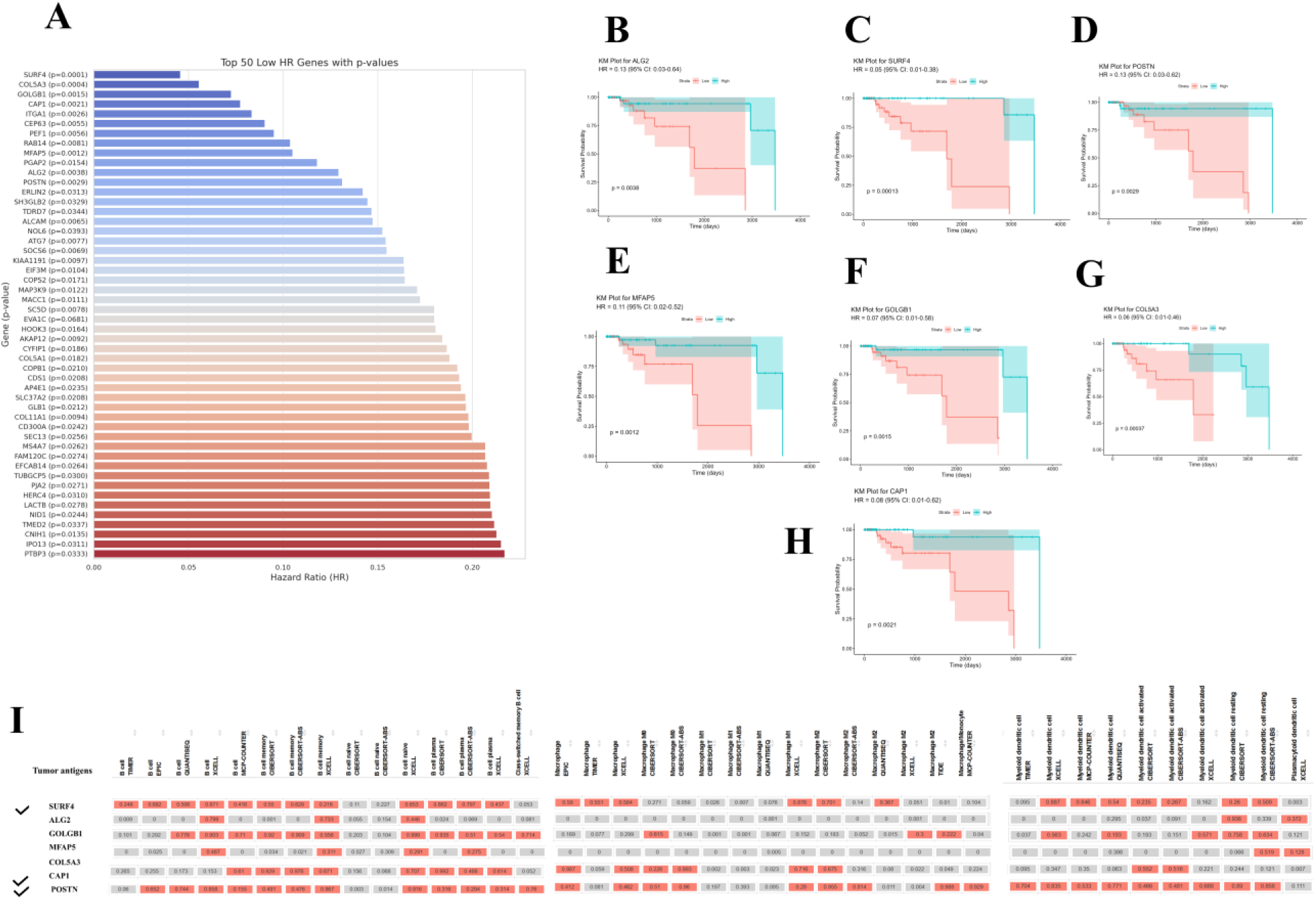
Overall survival analysis A) Forest plot showing p and HR values of top 50 tumor antigens, B-H) Overall survival plots of top 7 genes based on significance p-value (<0.005) and low HR value, I) Correlation of tumor antigens with APCs

**Figure 3.**
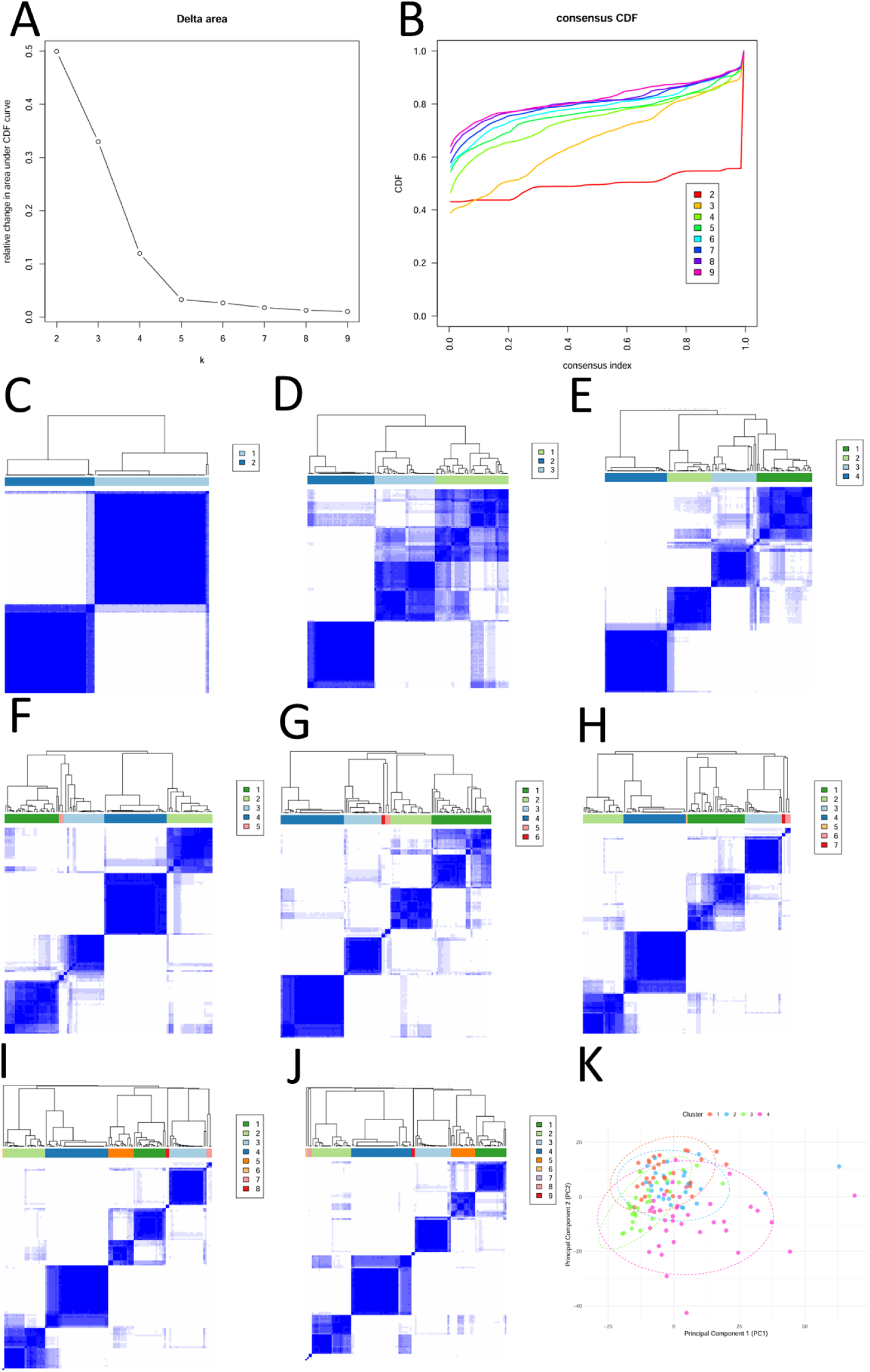
Consensus clustering of immune genes, A) Delta area, B) Cumulative distribution function (CDF) curve, C-J) clustering matrix of K from 2-9 clusters, K) PCA plot showing distribution of samples for k value 4

### 3.3. Analysis of Antigen-Immune Cell Correlations

TIMER analysis revealed significant correlations between the specific tumor antigens and the presence of antigen-presenting immune cells (B cells, dendritic cells, and macrophages) [51]. Among the seven tumor antigens analysed, three genes, *POSTN, SURF4*, and *CAP1*, exhibited strong correlations with these immune cell types. The correlations were determined based on a stringent p-value threshold of <0.005, ensuring the statistical relevance of the observed associations [52]. The correlation plots for these seven genes with B cells, dendritic cells, and macrophages are illustrated in Figure 2I, highlighting the connection between tumor antigen expression levels and the infiltration of immune cells within the tumor microenvironment. These results indicate that these tumor antigens may be actively involved in the immune response in TNBC, potentially influencing the effectiveness of immunotherapies [53].

### 3.4. Mutation incorporation in the screened tumour antigen to screen neoantigen peptides

The identified mutations in *POSTN, SURF4*, and *CAP1* span coding, intronic, and regulatory regions. In *POSTN*, two missense substitutions were detected in distinct samples: c.213G>T (p.Q71H) and c.46G>A (p.V16I), both localized to the coding region. *SURF4* carried two independent stop-gained variants within the same sample, c.349G>T (p.G117*) and c.478G>T (p.G160*), predicted to induce premature translational termination with potential loss of protein function. In *CAP1*, a missense mutation c.498C>G (p.N166K) was observed, resulting in the replacement of a neutral asparagine residue with a positively charged lysine within a critical cytoskeletal interaction domain. This substitution is predicted to perturb local charge distribution and compromise actin-dynamics–related functionality. To evaluate their immunotherapeutic applicability, each mutation was manually incorporated into the corresponding wild-type sequence for immunogenic neoantigen peptides. Since adaptive immune recognition is restricted to non-self peptide fragments, we focused on identifying mutant-derived epitopes. For each candidate, 15 amino acid windows were generated by extracting 9 residues upstream and downstream of the mutated position, with the mutation centered. This approach aligns with the binding preferences of both MHC class I and class II molecules, which typically accommodate peptides of 9–15 amino acids. From this screening, three putative neoantigenic peptides were prioritized: two derived from POSTN missense variants and one from the CAP1 substitution. The predicted sequences, along with their corresponding wild-type and mutant forms, are summarized in Table 1.

**Table 1.**
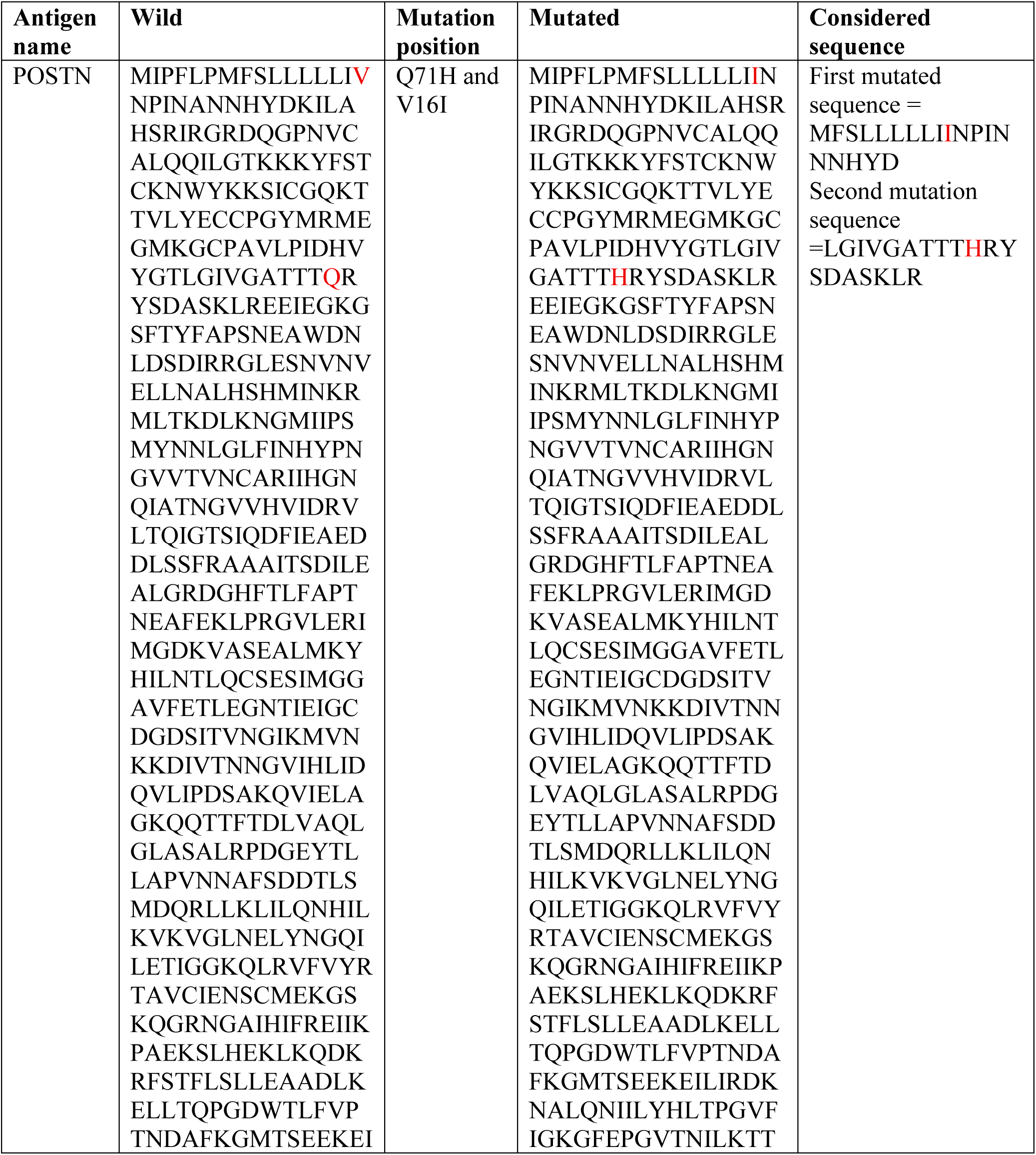
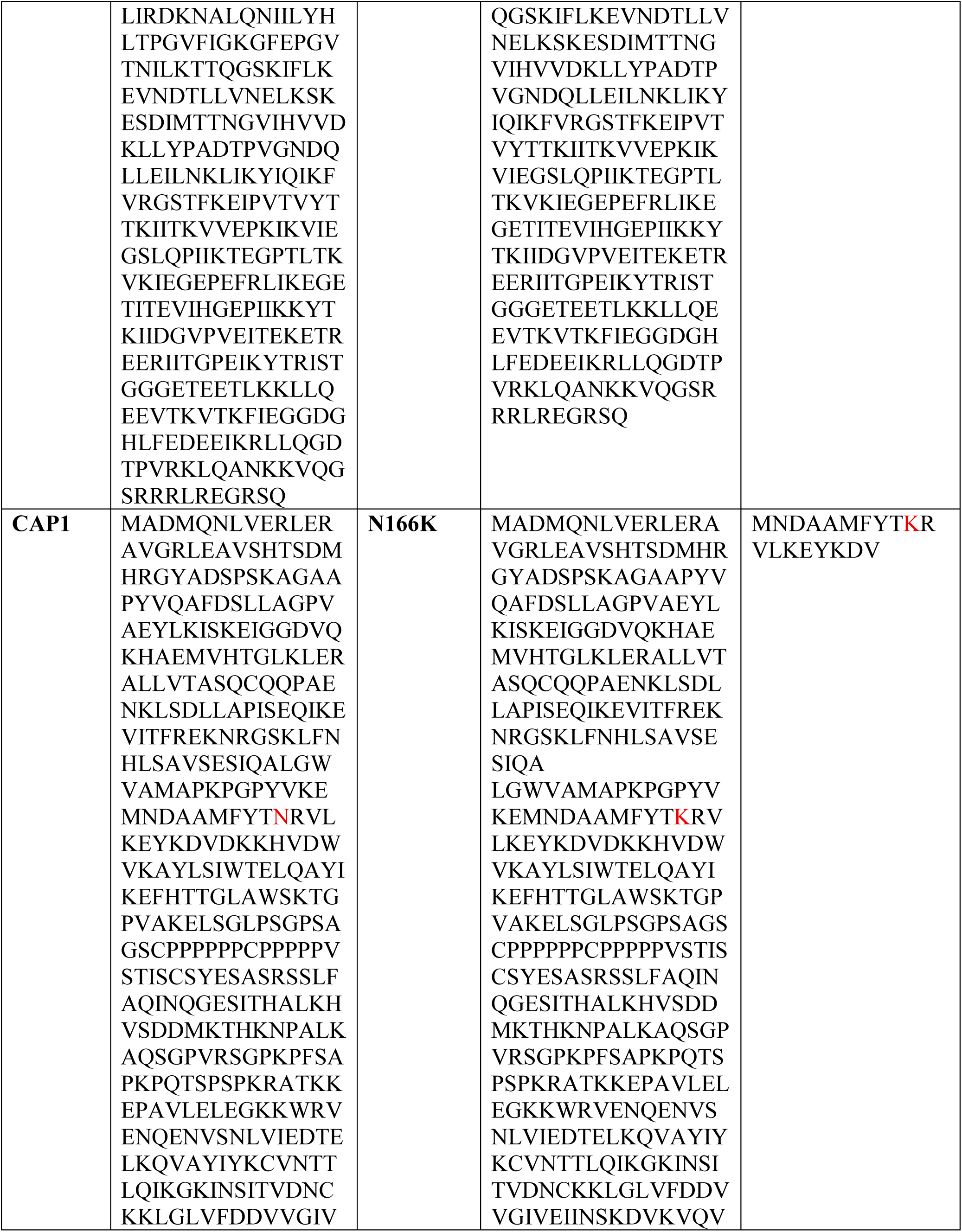
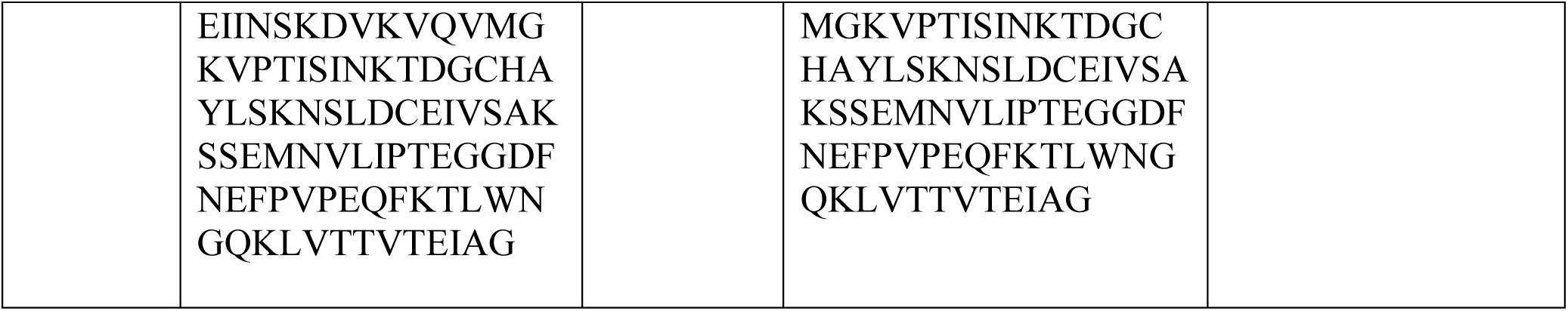
Neoantigen wild and mutated sequence.

### 3.5. Identification of the Immune Subtypes (IS) in TNBC

Consensus clustering was conducted on 121 TNBC samples, focusing on 1,753 immune-related genes. The delta area plot indicated a peak at a k value of 2, followed by a progressive decline up to a k value of 5, as illustrated in Figure 3A. The cumulative distribution function (CDF) curve demonstrated a smooth and stable profile for k value 4, suggesting consistent distribution during the clustering process (Figure 3B). In contrast, other k values displayed greater variability [54]. The consensus matrices for all nine k values are depicted in Figures 3C-J. Based on the CDF curve, four immune subtypes (IS) were identified: IS1 comprising 29 samples, IS2 with 26 samples, IS3 containing 29 samples, and IS4 being the largest, encompassing 37 samples. The distribution of these four immune subtypes is depicted in the PCA plot as shown in Figure 3K.

### 3.6. Mutational profile across Immune Subtypes

The oncoplot analysis reveals overlapping and distinct patterns of mutational profiles across the four immune subtypes of TNBC samples. As shown in Figure 4. In subtype 1 (Figure 4A), 96% (24/25) of samples exhibited mutations, with *TP53* being the gene with the highest mutation frequency (84%), followed by *MUC16* and *CSMD3* (12-16% mutation rates). Subtype 2 showed mutations in 91.67% (22/24) of samples, with *TP53* mutations present in 83% and other notable genes, including *CREBBP* and *AHNAK*, exhibiting mutation frequencies between 12% and 29% (Figure 4B). In subtype 3, all samples (100%) were mutated, highlighting *TP53* (76%) and *KMT2D* (20%) as the most commonly affected genes, with additional genes such as *MUC16, ATAD2B*, and *SYNE1* showing mutation rates of 12-16% (Figure 4C). Subtype 4 had 93.1% (27/29) of samples with mutations, where *TP53* was again the most frequently altered gene (79%), followed by *PIK3CA* (28%) and several other genes with mutation frequencies up to 14% (Figure 4D). *TP53* mutations are prevalent across all subtypes, indicating their common role in TNBC regardless of immune subtype. Additionally, mutational analysis revealed that immune subtype 1 had the lowest mutation burden of 152, the highest mutational burden was seen in IS 3 of 1138, but a high mutation burden was seen in just two samples, the remaining samples showed very low mutational burden. IS 2 showed 883, and IS4 of 737 [55–56].

**Figure 4.**
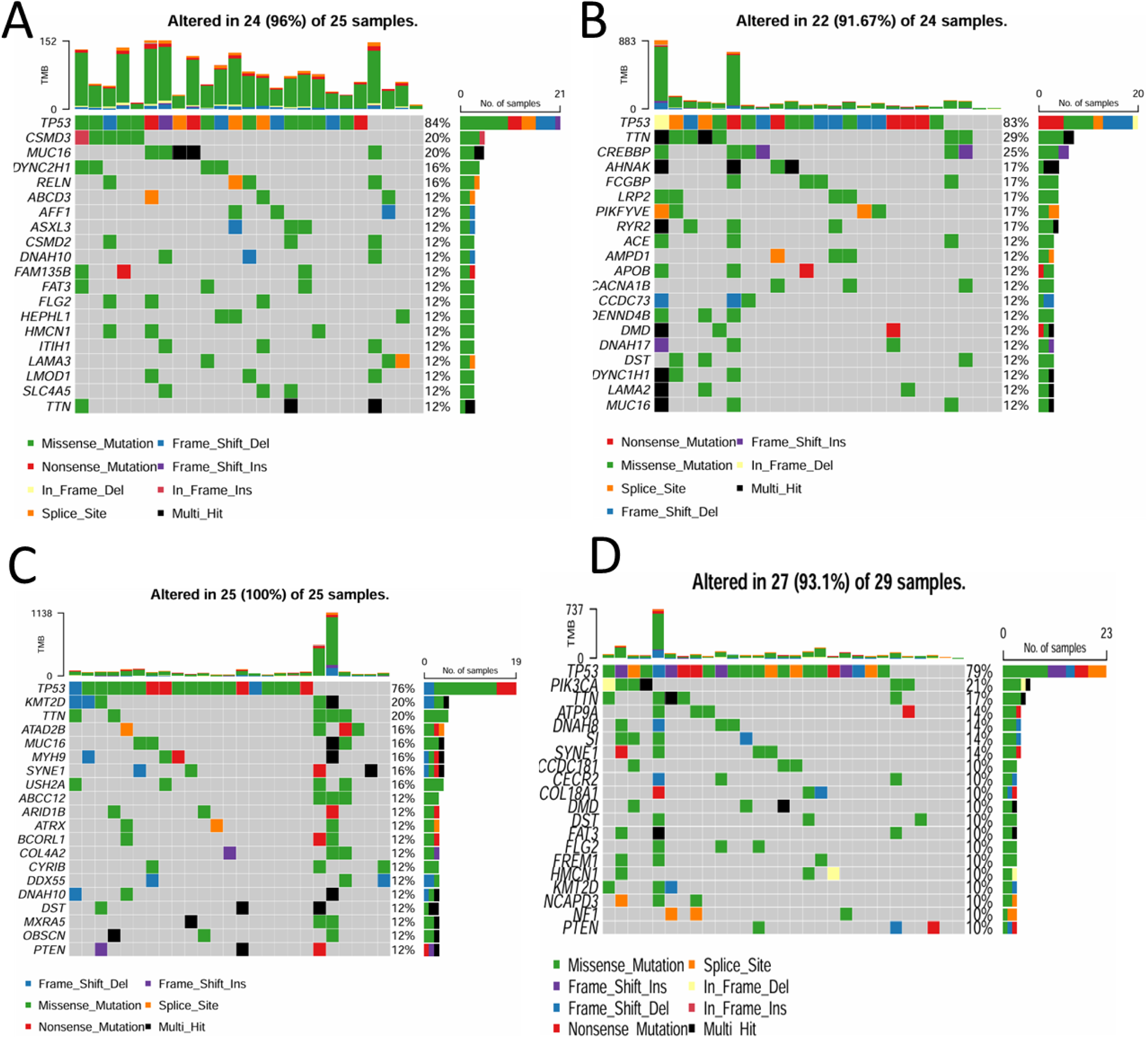
Mutational landscape of the top 20 mutated genes in all immune subtypes of TNBC, illustrated using Oncoplot (A-IS1, B-IS2, C-IS3, D-IS4)

### 3.7. Immune-related molecular features of immune subtypes

Figure 5 provides a comprehensive analysis of the four immune subtypes (IS1-4) in TNBC, focusing on their enrichment scores, immune cell composition, immune checkpoint gene expression, immune cell death genes, and overall survival outcomes. Figure 5A presents a heatmap illustrating the enrichment scores of 17 immune cell signatures across the subtypes. The heatmap analysis highlights distinct immune activity across the four subtypes. IS2 and IS4 stand out as the "immune-hot" subtype, with high enrichment in multiple immune functions like B cell and T cell activation and proliferation, indicating robust immune engagement. IS1 shows moderate enrichment in a few samples, and many samples had very low enrichment scores, suggesting an intermediate immune profile. IS3 exhibits a very low immune profile, representing an "immune-cold" subtype, with low enrichment across most immune functions, reflecting minimal immune activation or a more immunosuppressed state. Figure 5B provides the CIBERSORT heatmap. A very strong immune activity was seen in IS4, which represented a very strong immune-hot type, and IS3 showed very low immune activity, which makes them an immune-cold subtype, and IS1 and 2 showed moderate immune activity. ESTIMATE analysis results, showing stromal scores, immune scores, and combined estimate scores for each subtype. Each subtype exhibits a unique pattern in immune cell distribution. For instance, IS4 and IS2 consistently have higher immune, stromal, and estimate scores, and IS1 and IS3 have low scores, suggesting differences in the tumor microenvironment’s immune and stromal content across subtypes, as illustrated in Figure 5C-F.

**Figure 5.**
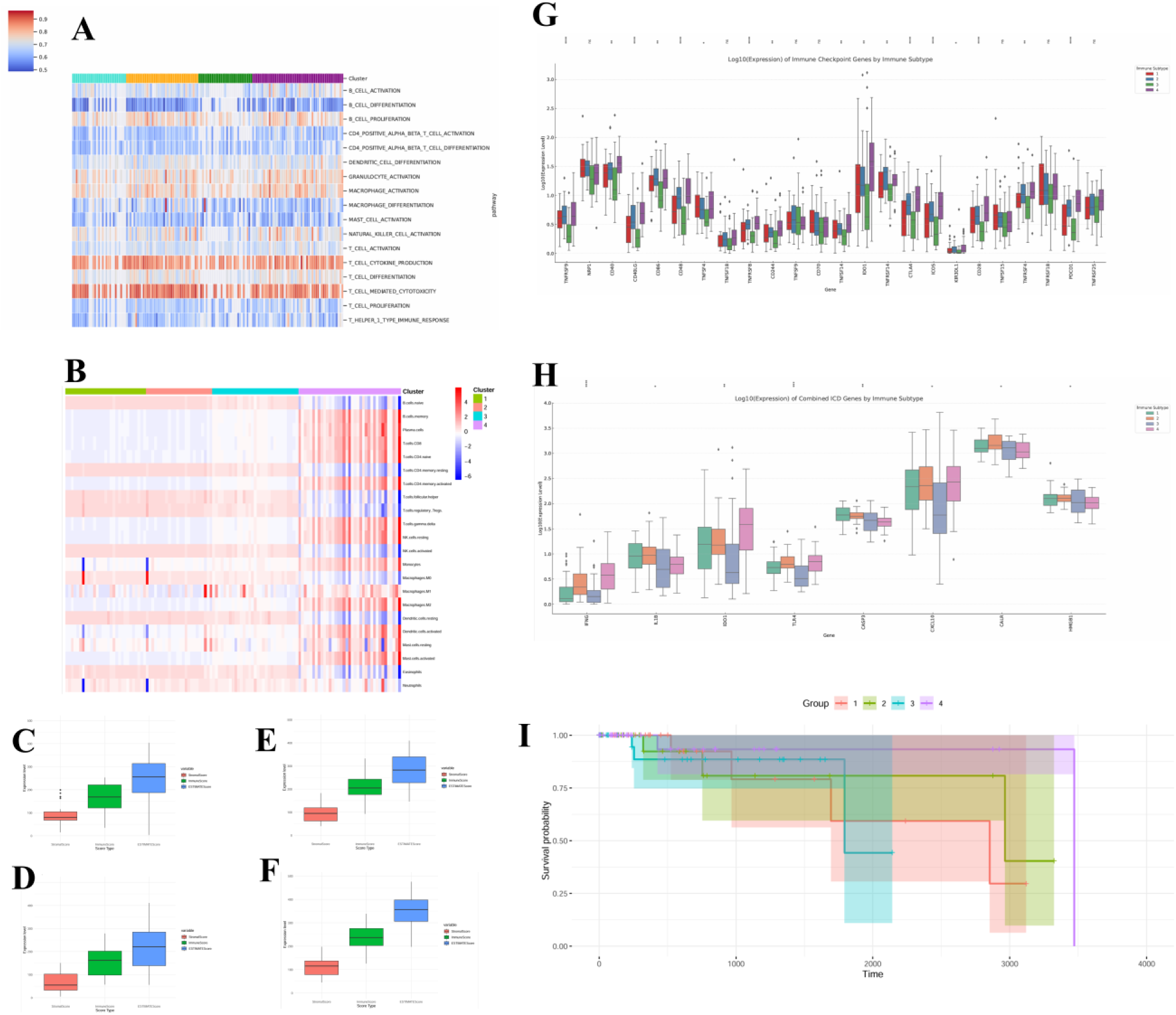
Immune profile analysis of four immune subtypes and their overall survival analysis A) Heatmap showing the enrichment scores of 17 immune cell signatures in TNBC, B) CIBERSORT profile showing the 22 immune cells activity in each IS, C-F) Estimate analysis boxplots for IS1-4, G) Immune checkpoint genes expression levels in each IS, H) Immune cell death genes of each IS, I) Overall survival analysis plots for each IS

### 3.8. Immune Checkpoint gene (ICP) and Immune Cell Death (ICD) genes profile in immune subtype and Prognostic analysis of immune subtypes

ICP genes regulate the immune system’s response, controlling the balance between excessive immune activity and autoimmunity, but in the context of cancer. ICP is produced by the cancer cells to suppress the immune cells. The presence of ICP directly represents the active immune response around the tumor cells, and ICD genes regulate various forms of programmed cell death (PCD) in immune cells, which is crucial for maintaining immune homeostasis, eliminating defective cells, and preventing excessive immune activation. These genes regulate processes such as apoptosis, pyroptosis, necroptosis, and ferroptosis in cancer cases. ICD genes are requested by dying cancer cells, thus triggering an active immune response against cancer cells. The presence of both ICP and ICD is more seen in IS 2 and 4 compared to IS 1 and 3 (Figure 5G and H), which suggests that ICP treatment is best suited for IS 2 and 4, blocking them would be beneficial from increased immune response, whereas for the IS 1 and 3 vaccine therapy would be beneficial combined with ICP. Figure 5I illustrates the overall survival analysis for the four subtypes. IS4, with its robust immune profile, shows relatively favourable survival outcomes. IS2 showed the second-best survival subtype, both of them have a very high immune-active status, which might have influenced the elevated OS rates. IS1 and IS3 showed bad progress characterized by low immune cell infiltration, minimal checkpoint gene expression, and poor immune cell death gene activity, which correlates with the least favourable survival, underscoring the need for strategies to boost immune activation in this subtype. IS1 and IS3 showed low mutational burden and released very low mutated neoantigens. Thus, immune response is very low in those groups, and externally treating the immune cell with an mRNA vaccine by incorporating neoantigen epitopes in the vaccine would activate immune response against those subtypes [57].

### 3.9. WGCNA to identify the biomarker to assess the effect of mRNA vaccine

WGCNA analysis was performed to identify co-expression modules among immune-related genes associated with the best survival rates in subtypes bearing a high immune profile. A soft-thresholding power of β = 6 was selected, yielding a scale-free topology with R² = 0.9 (Figures 6A, B). The representation matrix was converted into an adjacency matrix, which was then transformed into a topological overlap matrix. An average-linkage hierarchical clustering method was employed to analyze the data further to generate a dendrogram, applying a minimum module size criterion of 30 genes (Figure 6C). The gene dendrogram revealed 11 distinct modules, with the largest being the gray module, containing over 500+ genes, followed by the turquoise and yellow modules. Smaller black, blue, and green modules were also identified, each containing fewer than 100 genes (Figure 6D). A scatter plot of Eigengene Expression of 11 models in each immune subtype can be seen in Figure 6E, and a box plot of expression levels of identified model genes in each IS Can be seen in Figure 6F, in box plot each module showed high counts in IS 2 and IS4 with says immune cells count are more in IS2 and IS4 compare to IS 1 and 3 [58]

**Figure 6.**
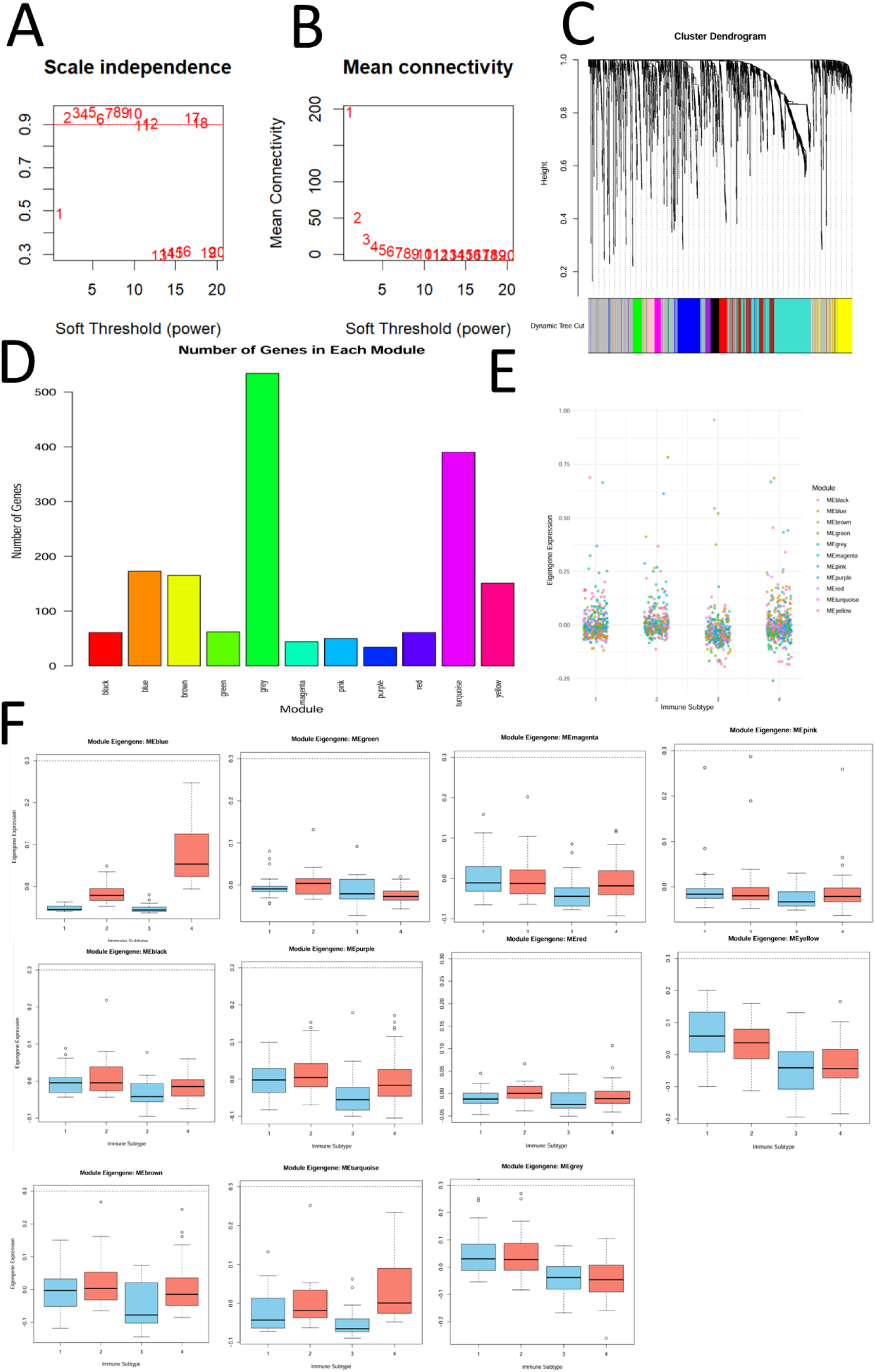
WGCNA analysis of immune genes in TNBC, A) The scale-free fit index across different soft-thresholding powers (β). B) The mean connectivity for various values of soft-thresholding powers., C) Dendrogram, D) Number of genes in each module, E) Scatter plot of Eigengene Expression in each IS, F) Expression levels of identified model genes in each IS

### 3.10. Hub Gene Identification in overall favourable modules

Prognosis analysis (Figure 7A) of these 11 models revealed better progression in gray, blue, and pink modules [59]. To know what pathways are involved in them, KEGG, biological function, and molecular function pathways were analysed, and in the pink module, pathways related to non-immune response were observed, but in blue and gray, many pathways were associated with the immune response, as shown in the bar plots of Figure 7B-J. Blue module exhibits more immune relevance, featuring KEGG pathways such as Th17 cell differentiation, cytokine-cytokine receptor interaction, and JAK-STAT signaling, all of which are involved in immune modulation. The biological processes in the blue module highlight immune-related functions like T cell activation and cytokine signaling, and its molecular functions indicate cytokine activity and receptor interactions, making it more immunologically relevant than the pink module. Similarly, the gray module also shows the strongest immune association, with KEGG pathways covering both innate and adaptive immune responses, including IL-17 signaling, Toll-like receptor signaling, T cell receptor signaling, natural killer (NK) cell-mediated cytotoxicity, and cytokine-cytokine receptor interaction. Its biological processes emphasize immune system activation, cytokine regulation, and cell activation, while its molecular functions include immune receptor activity and growth factor binding, indicating extensive immune involvement. Compared to the pink module, which is primarily developmental, the blue and gray module has the most immune-enriched, encompassing broad immune activation pathways, making it the most relevant in terms of immune response. Hub gene identification was performed for both the blue and gray modules, as both are associated with major immune response pathways. In the blue module, many hub genes are related to immunoglobulins, which contribute to the favorable overall response observed in this group. In the gray module, hub genes are more involved in cytokine responses, including IL-4, IL-6, and tumor necrosis factor genes, which are responsible for the favorable outcomes in this group, as illustrated in Figure 7K-N.

**Figure 7.**
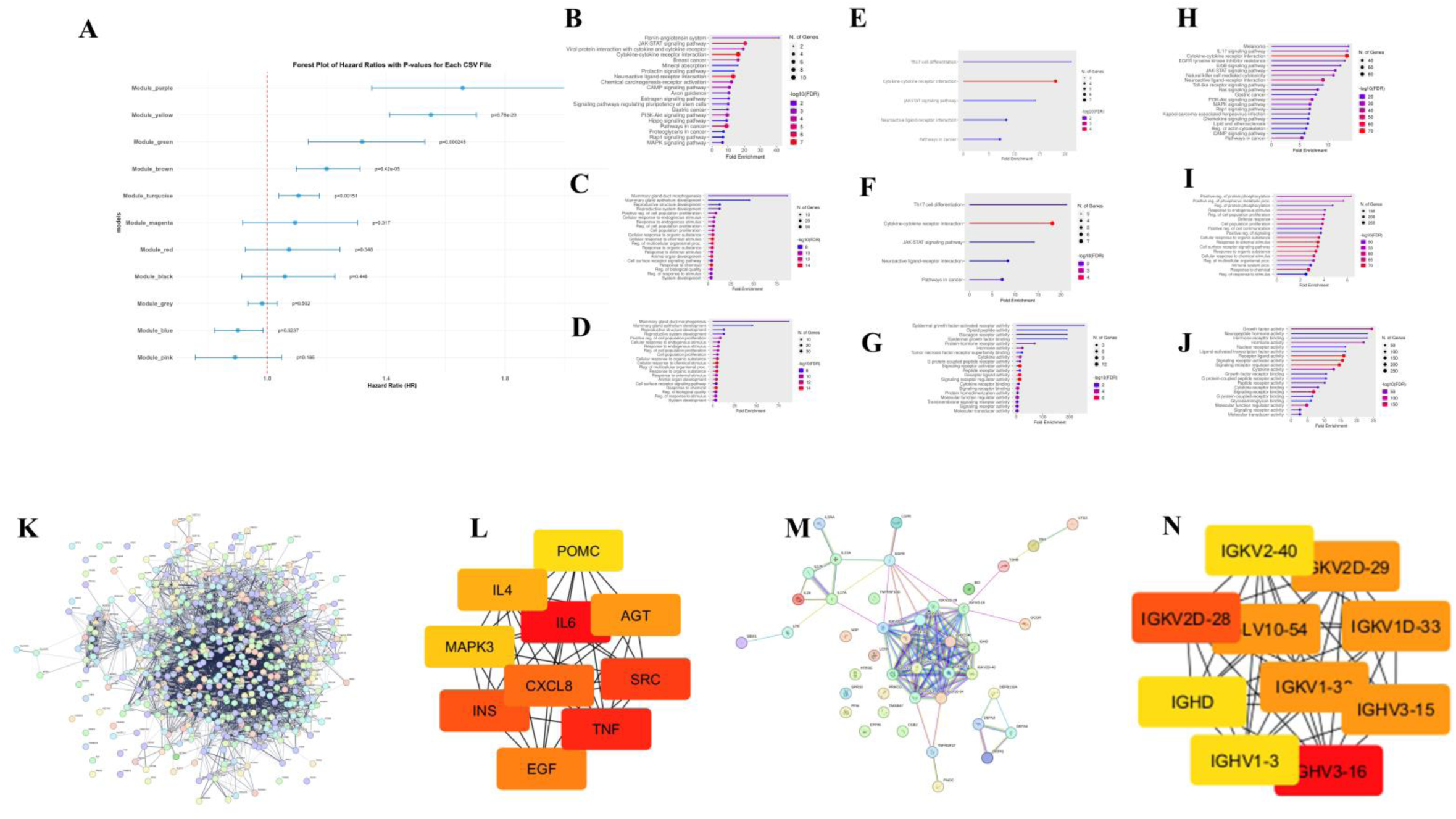
A) Prognosis analysis of all 11 model genes, B-D) KEGG pathway, biological process, and molecular function analysis of the pink model, E-F) KEGG pathway, biological process, and molecular function analysis of the blue model, H-J) KEGG pathway, biological process, and molecular function analysis of the gray module, K-L) PPI network and hub genes of the gray module, M-NPPI network and hub genes of the blue module.

## 4. Discussion

TNBC is a highly aggressive breast cancer subtype that is difficult to treat because it lacks ER, PR, and HER2 [1]. As a result, targeted therapies such as hormonal treatments or HER2 inhibitors, commonly utilized in other breast cancer subtypes, are ineffective against TNBC [4]. Consequently, chemotherapy remains the primary treatment, but the aggressive nature of TNBC with a high level of recurrence and metastasis leads to poor survival outcomes [2, 6]. In recent years, immunotherapy, specifically immune checkpoint inhibitors (ICIs), has emerged as a potential treatment option for TNBC. However, the overall response rate remains limited, with only a small proportion of patients deriving benefits from these therapies [5]. This highlights a critical gap in the development of effective treatments for TNBC, particularly therapies that can be tailored to target the molecular and immunological characteristics of the disease. There is an urgent demand for innovative approaches, such as cancer vaccines, which can activate the immune system to target tumor cells [19]. In light of these challenges, this study aimed to identify novel tumor-neoantigens and screen populations suitable for vaccination, followed by biomarker identification to monitor patent’s progress.

Integration of differential gene expression analysis and mutation profiling with CNA genes identified 2264 genes that were both significantly upregulated and mutated with amplified CNA in TNBC samples, making them potential tumor antigens. These antigens may act as key candidates for mRNA vaccine designs because they are tumor-specific, which is crucial for reducing off-target effects and increasing immune specificity. The prognosis evaluation highlighted seven genes (*ALG2, SURF4, CAP1, COL5A3, POSTN, GOLGB1*, and *MFAP5*) that are significantly correlated with patient prognosis. The KM plot and HR values for these genes suggest that individuals with high expression levels of these genes have a lower risk of recurrence and a better prognosis [60]. For these seven tumor antigens, correlation analysis with antigen-presenting cells (APCs), including B cells, dendritic cells, and macrophages, was performed to examine their role in immune recognition. APCs play a vital role in initiating and modulating the immune response by processing tumor antigens and presenting their fragments to T cells, which facilitates cancer cell recognition and destruction. Effective presentation of tumor antigens by APCs is essential for activating cytotoxic T cells that target and eliminate tumor cells [61]. The correlation analysis between these tumor antigens and APC prevalence provided valuable insights into their immunogenicity and potential to elicit an immune response in TNBC. Among the seven identified tumor antigens, three genes *(POSTN, SURF 4*, and *CAP 1*) showed strong correlations with APCs. However, in *SURF 4,* premature proteins are being translated, so we did not include it for the neoantigen peptide selection. Finally, by manual incorporation of mutations in them, we present 3 neoantigen peptides suitable for vaccine design. These may be actively recognized by APCs and play a key role in promoting antigen presentation and subsequent T cell activation, making them promising vaccine candidates.

To determine which patient groups would benefit most from an mRNA vaccine, immune subtyping was performed to assess the immune profile. This analysis helps understand the distribution and activity of immune cells, mutation burden in each subtype, ICP, and immunogenic cell death (ICD), and their influence on patents survival rates in TNBC patients, providing insights into immune cell infiltration patterns within the tumor microenvironment.

Subtyping revealed significant heterogeneity among tumor samples. Using consensus clustering, four distinct immune subtypes were identified, each characterized by unique patterns of immune cell infiltration and mutational profiles. This diversity in immune infiltration is crucial for understanding how different TNBC subtypes may respond to immunotherapy. The mutational landscape analysis across the four immune subtypes of TNBC provides critical insights into their mutational burden and its relation with immunological profiles. The results reveal the ubiquity of *TP53* mutations, which is a tumor suppressor gene across all subtypes, strengthening its role as a common driver in TNBC regardless of immune characteristics. This analysis revealed that the mutation of the *TP53* gene may be the reason for the cancer cells arising. Additionally, the mutation burden in each group revealed unique patterns. IS 1 and 3 showed very low mutational burden compared to IS 2 and 4, and due to decreased mutations, very low neoantigens may be produced, and low immune response may be seen [62–64].

To confirm the relation between mutational burden and immune response, immune profiles across the four IS were analyzed. IS2 and IS4 demonstrate substantial immune cell infiltration, particularly of cytotoxic T cells, which are crucial for an effective immune response against tumors. The notable enrichment of cytotoxic T cells, CD8+ T cells, B cells, macrophages, and various other immune pathways in these immune subtypes suggests a "hot" tumor microenvironment. CIBERSORT analysis also showed IC3 as a very weak immune profile group [65]. Notably, IS2 and 4 are characterized by high levels of ICD and ICP, indicating that these tumors may be more vulnerable to ICIs in combination with other treatments, such as mRNA vaccines [66–67]. The high expression of immune checkpoints in IS 2 and 4 further reinforces the presence of immune cells around the tumor, but due to the ICP genes, tumor neutralization is not seen. IS 1 and 3 display low levels of immune infiltration and immune activity compared to IS 2 and 4. IS 3, in particular, shows the lowest immune scores, suggesting that these tumors have a "cold" immune microenvironment [68].

The ESTIMATE analysis provided further insights into the immune landscape by calculating stromal, immune, and overall ESTIMATE scores, which reflect the distribution of immune and stromal cells within the tumor microenvironment. IS 4 exhibits the highest stromal and immune scores, indicating elevated levels of immune cell infiltration and stromal activity. This suggests that these IS have a robust immune presence. IS 1 and 3 show moderate immune and stromal scores. Further, upon survival analysis of the subtypes, it was observed that higher survival rates were observed in IS 4 and 2, and it was clear that the immune profile is influencing the survival rates [69].

Based on this analysis, externally recognizing immune cells with tumor antigens could enhance the immune response in IS 1 and 3, potentially converting their "cold" immune microenvironment into a more immunogenic state. Given that IS 2 and 4 exhibit high immune infiltration and a favorable response to immune checkpoint inhibitors (ICIs), a strategic approach to boost the immune activity in IS 1 and 3 would be to facilitate external immune recognition. One promising method to achieve this is through the design of an epitope-based vaccine using an mRNA vaccine. mRNA vaccines can be engineered to encode epitopes of these neoantigens. By incorporating highly immunogenic epitopes, these vaccines can promoting a robust anti-tumor immune response. Given the observed low mutational burden and reduced neoantigen production in these subtypes, designing vaccines that induce presentation of tumor neoantigens epitopes could help bypass immune evasion mechanisms. Ultimately, leveraging mRNA-based epitope vaccines tailored to the immune-deficient characteristics of IS 1 and 3 could transform their immune landscape, making them more responsive to immunotherapy. This approach holds significant potential in improving TNBC cases that currently exhibit low immune activity and resistance to conventional treatments.

To further investigate biomarker genes that help in monitoring the immune response post vaccination, WGCNA analysis was carried out to screen genes that are responsible for favorable survival rates in IS 2 and 4. Thus, post vaccination, monitoring these genes in patients having a similar immune profile to IS1 and 3 would help in evaluating patents progress. Among these, the gray module, pink module, and blue module showed the best survival rates, and pathway analyses revealed that the blue and gray modules showed more immune-related pathways in them, and hub genes were identified for these modules and revealed the genes coding for cytolines like IL 6 and 4 TNF gens and immunoglobulin genes were responsible for favourable survival status of IS 2 and 4 groups [70–73].

## Conclusion

This study comprehensively analyses the immune landscape in TNBC and highlights the potential for developing targeted immunotherapies, particularly mRNA vaccines. By identifying novel tumor antigens such as *POSTN* and *CAP1*, which are strongly associated with antigen-presenting cells, we have outlined promising candidates for immune activation. Our immune subtyping revealed distinct immune profiles across TNBC subtypes, with IS 1 and IS 3 emerging as the most favourable candidates for immunotherapy due to their low immune infiltration and checkpoint expression.

## Supporting information

https://docs.google.com/document/d/14SeldeYgglvlhysnPiHOUWhPzmMm5xCi/edit?usp=drive_link&ouid=114036230192474557622&rtpof=true&sd=true

## Authorship contribution statement

**Karthick Vasudevan:** Writing – review & editing, Project administration, Supervision, Conceptualization, Visualization, Validation, Methodology, Formal analysis, Data curation. **Dhanushkumar T:** Writing – review & editing, Writing – original draft, Visualization, Validation, Methodology, Formal analysis, Data curation, Conceptualization. **Prasanna Kumar Selvam:** Writing – review & editing, Writing – original draft, Visualization, Formal analysis, Data curation. **Anbarasu Krishnan:** Writing – review & editing, Writing – original draft. **Sunila B G:** Writing – review & editing, Writing – original draft, Methodology. **Santhosh Mudipalli Elavarase:** Review & editing, validation. **Supraja Mohan:** Writing – review & editing, Writing – original draft. **Rohini Karunakaran:** Writing – review & editing, Project administration, Supervision, Conceptualization, Visualization, Validation, Methodology, Formal analysis, Data curation.

## Conflict of interest

The authors declare no conflict of interest.

## Funding

No funding was received for conducting this study.

## Data availability

All data generated or analyzed during this study are included in this article and supplementary materials.

## Limitations and future direction

One of the main limitations of this study is the lack of validation of the findings in both *in vitro/in vivo* conditions. However, the computational findings provide a strong foundation for using the identified mutated peptides as a source for epitope prediction and developing an mRNA vaccine. Additionally, the data source used for these findings is from a single source (TCGA). Therefore, future directions should focus on to cross -validating the mutations in the data from additional independent datasets and designing a vaccine from the epitope predictions from the identified mutated peptides. We strongly recommend using our cancer-specific epitope prediction tool from our previous study (VaxOptiML and DeepEpitope) for this purpose, as it is completely trained on cancer-specific epitope datasets using advanced machine learning and deep learning models, ensuring higher prediction accuracy and biological relevance.

## Notes

### Competing Interest Statement

The authors have declared no competing interest.

## References

1. Gucalp A, Traina TA. Triple-negative breast cancer: role of the androgen receptor. Cancer J. 2010;16(1):62–5.

2. Aysola K, Desai A, Welch C, Xu J, Qin Y, Reddy V, et al. Triple negative breast cancer–an overview. Hered Genet Curr Res. 2013;2013(Suppl 2).

3. de Ruijter TC, Veeck J, de Hoon JP, van Engeland M, Tjan-Heijnen VC. Characteristics of triple-negative breast cancer. J Cancer Res Clin Oncol. 2011;137:183–92.

4. Kirkby M, Popatia AM, Lavoie JR, Wang L. The potential of hormonal therapies for treatment of triple-negative breast cancer. Cancers (Basel). 2023;15(19):4702.

5. Farshbafnadi M, Khoshbin AP, Rezaei N. Immune checkpoint inhibitors for triple-negative breast cancer: from immunological mechanisms to clinical evidence. Int Immunopharmacol. 2021;98:107876.

6. Kumar S, Chatterjee M, Ghosh P, Ganguly KK, Basu M, Ghosh MK. Targeting PD-1/PD-L1 in cancer immunotherapy: an effective strategy for treatment of triple-negative breast cancer (TNBC) patients. Genes Dis. 2023;10(4):1318–50.

7. Zhu X, Zhang Q, Wang D, Liu C, Han B, Yang JM. Expression of PD-L1 attenuates the positive impacts of high-level tumor-infiltrating lymphocytes on prognosis of triple-negative breast cancer. Cancer Biol Ther. 2019;20(8):1105–12.

8. Sun WY, Lee YK, Koo JS. Expression of PD-L1 in triple-negative breast cancer based on different immunohistochemical antibodies. J Transl Med. 2016;14:1–12.

9. Schuster M, Nechansky A, Kircheis R. Cancer immunotherapy. Biotechnol J. 2006;1(2):138–47.

10. Stanculeanu DL, Daniela Z, Lazescu A, Bunghez R, Anghel R. Development of new immunotherapy treatments in different cancer types. J Med Life. 2016;9(3):240.

11. Vergati M, Intrivici C, Huen NY, Schlom J, Tsang KY. Strategies for cancer vaccine development. Biomed Res Int. 2010;2010(1):596432.

12. Hirayama M, Nishimura Y. The present status and future prospects of peptide-based cancer vaccines. Int Immunol. 2016;28(7):319–28.

13. Aurisicchio L, Ciliberto G. Genetic cancer vaccines: current status and perspectives. Expert Opin Biol Ther. 2012;12(8):1043–58.

14. Santos PM, Butterfield LH. Dendritic cell–based cancer vaccines. J Immunol. 2018;200(2):443–9.

15. Mittendorf EA, Ardavanis A, Symanowski J, Murray JL, Shumway NM, Litton JK, et al. Primary analysis of a prospective, randomized, single-blinded phase II trial evaluating the HER2 peptide AE37 vaccine in breast cancer patients to prevent recurrence. Ann Oncol. 2016;27(7):1241–8.

16. Norton N, Youssef B, Hillman DW, Nassar A, Geiger XJ, Necela BM, et al. Folate receptor alpha expression associates with improved disease-free survival in triple negative breast cancer patients. NPJ Breast Cancer. 2020;6(1):4.

17. Zhang X, Goedegebuure SP, Myers NB, Vickery T, McLellan MD, Gao F, et al. Neoantigen DNA vaccines are safe, feasible, and capable of inducing neoantigen-specific immune responses in patients with triple negative breast cancer. medRxiv. 2021 Nov.

18. Benvenuto M, Focaccetti C, Izzi V, Masuelli L, Modesti A, Bei R. Tumor antigens heterogeneity and immune response-targeting neoantigens in breast cancer. Semin Cancer Biol. 2021;72:65–75.

19. Lorentzen CL, Haanen JB, Met Ö, Svane IM. Clinical advances and ongoing trials of mRNA vaccines for cancer treatment. Lancet Oncol. 2022;23(10):e450–8.

20. Vishweshwaraiah YL, Dokholyan NV. mRNA vaccines for cancer immunotherapy. Front Immunol. 2022;13:1029069.

21. Miao L, Zhang Y, Huang L. mRNA vaccine for cancer immunotherapy. Mol Cancer. 2021;20(1):41.

22. Buonaguro L, Petrizzo A, Tornesello ML, Buonaguro FM. Translating tumor antigens into cancer vaccines. Clin Vaccine Immunol. 2011;18(1):23–34.

23. Tomczak K, Czerwińska P, Wiznerowicz M. The Cancer Genome Atlas (TCGA): an immeasurable source of knowledge. Contemp Oncol (Pozn). 2015;2015(1):68–77.

24. Mayakonda A, Koeffler HP. Maftools: Efficient analysis, visualization and summarization of MAF files from large-scale cohort based cancer studies. bioRxiv. 2016;052662.

25. Liu S, Wang Z, Zhu R, Wang F, Cheng Y, Liu Y. Three differential expression analysis methods for RNA sequencing: limma, EdgeR, DESeq2. J Vis Exp. 2021;(175):e62528.

26. Mayakonda A, Lin DC, Assenov Y, Plass C, Koeffler HP. Maftools: efficient and comprehensive analysis of somatic variants in cancer. Genome Res. 2018;28(11):1747–56.

27. Adamu PI, Adamu MO, Okagbue HI, Opoola L, Bishop SA. Survival analysis of cancer patients in north eastern Nigeria from 2004–2017–a Kaplan-Meier method. Open Access Maced J Med Sci. 2019;7(4):643.

28. Pawar A, Chowdhury OR, Salvi O. A narrative review of survival analysis in oncology using R. Cancer Res Stat Treat. 2022;5(3):554–61.

29. Nagy Á, Munkácsy G, Győrffy B. Pancancer survival analysis of cancer hallmark genes. Sci Rep. 2021;11(1):6047.

30. Li T, Fu J, Zeng Z, Cohen D, Li J, Chen Q, et al. TIMER2.0 for analysis of tumor-infiltrating immune cells. Nucleic Acids Res. 2020;48(W1):W509–14.

31. Bhattacharya S, Andorf S, Gomes L, Dunn P, Schaefer H, Pontius J, et al. ImmPort: disseminating data to the public for the future of immunology. Immunol Res. 2014;58:234–9.

32. Wilkerson M, Waltman P, Wilkerson MM. Package ‘ConsensusClusterPlus’. 2013. [Software Package].

33. Subramanian A, Kuehn H, Gould J, Tamayo P, Mesirov JP. GSEA-P: a desktop application for Gene Set Enrichment Analysis. Bioinformatics. 2007;23(23):3251–3.

34. Korotkevich G, Sukhov V, Budin N, Shpak B, Artyomov MN, Sergushichev A. Fast gene set enrichment analysis. bioRxiv. 2016;060012

35. Yoshihara K, Shahmoradgoli M, Martínez E, Vegesna R, Kim H, Torres-Garcia W, et al. Inferring tumour purity and stromal and immune cell admixture from expression data. Nat Commun. 2013;4(1):2612.

36. Chen B, Khodadoust MS, Liu CL, Newman AM, Alizadeh AA. Profiling tumor infiltrating immune cells with CIBERSORT. In: Cancer Systems Biology: Methods and Protocols. New York: Humana Press; 2018. p. 243–59.

37. Therneau T, Lumley T. R survival package. R Core Team. 2013;523.

38. Langfelder P, Horvath S. WGCNA: an R package for weighted correlation network analysis. BMC Bioinformatics. 2008;9:559.

39. Dawath B, Patole H, Dsouza N, Purohit K, Peter S. Identification of hub genes in the fructose-1,6-bisphosphate 1-protein interaction network based on differential expression, and survival impact on hepatocellular carcinoma patients. Biomed Biotechnol Res J. 2023;7(2):238–46.

40. Ge SX, Jung D, Yao R. ShinyGO: a graphical gene-set enrichment tool for animals and plants. Bioinformatics. 2020;36(8):2628–9.

41. von Mering C, Huynen M, Jaeggi D, Schmidt S, Bork P, Snel B. STRING: a database of predicted functional associations between proteins. Nucleic Acids Res. 2003 Jan 1;31(1):258–61.

42. Shannon P, Markiel A, Ozier O, Baliga NS, Wang JT, Ramage D, et al. Cytoscape: a software environment for integrated models of biomolecular interaction networks. Genome Res. 2003 Nov;13(11):2498–504.

43. Zhao D, Liu X, Shan Y, Li J, Cui W, Wang J, et al. Recognition of immune-related tumor antigens and immune subtypes for mRNA vaccine development in lung adenocarcinoma. Comput Struct Biotechnol J. 2022;20:5001–13.

44. Li R, He Y, Zhang H, Wang J, Liu X, Liu H, et al. Identification and validation of immune molecular subtypes in pancreatic ductal adenocarcinoma: implications for prognosis and immunotherapy. Front Immunol. 2021;12:690056.

45. Liu S, Wang Z, Zhu R, Wang F, Cheng Y, Liu Y. Three differential expression analysis methods for RNA sequencing: limma, EdgeR, DESeq2. J Vis Exp. 2021;(175):e62528.

46. Ye L, Wang L, Yang JA, Hu P, Zhang C, Tong SA, et al. Identification of tumor antigens and immune subtypes in lower grade gliomas for mRNA vaccine development. J Transl Med. 2021;19:1–13.

47. Li R, He Y, Zhang H, Wang J, Liu X, Liu H, et al. Identification and validation of immune molecular subtypes in pancreatic ductal adenocarcinoma: implications for prognosis and immunotherapy. Front Immunol. 2021;12:690056.

48. Xu H, Zheng X, Zhang S, Yi X, Zhang T, Wei Q, et al. Tumor antigens and immune subtypes guided mRNA vaccine development for kidney renal clear cell carcinoma. Mol Cancer. 2021;20:1–7.

49. Liang F, Zhang S, Wang Q, Li W. Treatment effects measured by restricted mean survival time in trials of immune checkpoint inhibitors for cancer. Ann Oncol. 2018;29(5):1320–4.

50. Wei Y, Zheng L, Yang X, Luo Y, Yi C, Gou H. Identification of immune subtypes and candidate mRNA vaccine antigens in small cell lung cancer. Oncologist. 2023;28(11):e1052–64.

51. Jiang J, Pan W, Xu Y, Ni C, Xue D, Chen Z, et al. Tumour-infiltrating immune cell-based subtyping and signature gene analysis in breast cancer based on gene expression profiles. J Cancer. 2020;11(6):1568.

52. Tan H, Yu T, Liu C, Wang Y, Jing F, Ding Z, et al. Identifying tumor antigens and immuno-subtyping in colon adenocarcinoma to facilitate the development of mRNA vaccine. Cancer Med. 2022;11(23):4656–72.

53. Gao G, Fang M, Xu P, Chen B. Identification of three immune molecular subtypes associated with immune profiles, immune checkpoints, and clinical outcome in multiple myeloma. Cancer Med. 2021;10(20):7395–403.

54. Olivier M, Hollstein M, Hainaut P. TP53 mutations in human cancers: origins, consequences, and clinical use. Cold Spring Harb Perspect Biol. 2010;2(1):a001008.

55. Faundes V, Malone G, Newman WG, Banka S. A comparative analysis of KMT2D missense variants in Kabuki syndrome, cancers and the general population. J Hum Genet. 2019;64(2):161–70.

56. Dufresne A, Lesluyes T, Ménétrier-Caux C, Brahmi M, Darbo E, Toulmonde M, et al. Specific immune landscapes and immune checkpoint expressions in histotypes and molecular subtypes of sarcoma. Oncoimmunology. 2020;9(1):1792036.

57. Cao D, Chen MK, Zhang QF, Zhou YF, Zhang MY, Mai SJ, et al. Identification of immunological subtypes of hepatocellular carcinoma with expression profiling of immune-modulating genes. Aging (Albany NY). 2020;12(12):12187.

58. Tian Z, He W, Tang J, Liao X, Yang Q, Wu Y, et al. Identification of important modules and biomarkers in breast cancer based on WGCNA. Onco Targets Ther. 2020;13:6805–17.

59. Qayoom H, Alkhanani M, Almilaibary A, Alsagaby SA, Mir MA. Mechanistic elucidation of Juglanthraquinone C targeting breast cancer: a network pharmacology-based investigation. Saudi J Biol Sci. 2023;30(7):103705.

60. Sabzehali F, Rahimi H, Goudarzi H, Goudarzi M, Izad MHY, Chirani AS, et al. Functional engineering of OprF-OprI-PopB as a chimeric immunogen and its cross-protective evaluation with GM-CSF against Pseudomonas aeruginosa: a comprehensive immunoinformatics evaluation. Inform Med Unlocked. 2021;25:100673.

61. Soleymani S, Janati-Fard F, Housaindokht MR. Designing a bioadjuvant candidate vaccine targeting infectious bursal disease virus (IBDV) using viral VP2 fusion and chicken IL-2 antigenic epitope: a bioinformatics approach. Comput Biol Med. 2023;163:107087.

62. Henry F, Boisteau O, Bretaudeau L, Lieubeau B, Meflah K, Grégoire M. Antigen-presenting cells that phagocytose apoptotic tumor-derived cells are potent tumor vaccines. Cancer Res. 1999;59(14):3329–32.

63. Shi Y, Jin J, Ji W, Guan X. Therapeutic landscape in mutational triple negative breast cancer. Mol Cancer. 2018;17:1–11.

64. Zhou S, Liu S, Zhao L, Sun HX. A comprehensive survey of genomic mutations in breast cancer reveals recurrent neoantigens as potential therapeutic targets. Front Oncol. 2022;12:786438.

65. Wang L, Geng H, Liu Y, Liu L, Chen Y, Wu F, et al. Hot and cold tumors: immunological features and the therapeutic strategies. MedComm. 2023;4(5):e343.

66. Ahmed A, Tait SW. Targeting immunogenic cell death in cancer. Mol Oncol. 2020;14(12):2994–3006.

67. Darvin P, Toor SM, Sasidharan Nair V, Elkord E. Immune checkpoint inhibitors: recent progress and potential biomarkers. Exp Mol Med. 2018;50(12):1–11.

68. Collins JM, Redman JM, Gulley JL. Combining vaccines and immune checkpoint inhibitors to prime, expand, and facilitate effective tumor immunotherapy. Expert Rev Vaccines. 2018;17(8):697–705.

69. Zhao D, Liu X, Shan Y, Li J, Cui W, Wang J, et al. Recognition of immune-related tumor antigens and immune subtypes for mRNA vaccine development in lung adenocarcinoma. Comput Struct Biotechnol J. 2022;20:5001–13.

70. Xu H, Zheng X, Zhang S, Yi X, Zhang T, Wei Q, et al. Tumor antigens and immune subtypes guided mRNA vaccine development for kidney renal clear cell carcinoma. Mol Cancer. 2021;20:1–7.

71. Ye L, Wang L, Yang JA, Hu P, Zhang C, Tong SA, et al. Identification of tumor antigens and immune subtypes in lower grade gliomas for mRNA vaccine development. J Transl Med. 2021;19:1–13.

72. Weng W, Yu L, Li Z, Tan C, Lv J, Lao IW, et al. The immune subtypes and landscape of sarcomas. BMC Immunol. 2022;23(1):46.

73. Wei Y, Zheng L, Yang X, Luo Y, Yi C, Gou H. Identification of immune subtypes and candidate mRNA vaccine antigens in small cell lung cancer. Oncologist. 2023;28(11):e1052–64.

