## Supplementary material for "Transcriptomics and mutational analysis to screen immunogenic neoantigen peptides and Patient stratification based on immune subtypes for TNBC": https://docs.google.com/document/d/14SeldeYgglvlhysnPiHOUWhPzmMm5xCi/edit?usp=drive_link&ouid=114036230192474557622&rtpof=true&sd=true

**Supplementary tables**

Table 1. Clinical information of patients used in this study

| Sample_ID | LastFollowupDays | DeathDays | VitalStatus |
| --- | --- | --- | --- |
| TCGA-A1-A0SK | NA | 967 | Dead |
| TCGA-A1-A0SP | 584 | NA | Alive |
| TCGA-A2-A04U | 671 | NA | Alive |
| TCGA-A2-A0CM | NA | 754 | Dead |
| TCGA-A2-A0D0 | 643 | NA | Alive |
| TCGA-A2-A0D2 | 761 | NA | Alive |
| TCGA-A2-A0SX | 1288 | NA | Alive |
| TCGA-A2-A0T0 | 337 | NA | Alive |
| TCGA-A2-A0T2 | NA | 255 | Dead |
| TCGA-A2-A0YE | 253 | NA | Alive |
| TCGA-A2-A1G6 | 132 | NA | Alive |
| TCGA-A2-A25F | 114 | NA | Alive |
| TCGA-A2-A3XT | 2238 | NA | Alive |
| TCGA-A2-A3XX | 1168 | NA | Alive |
| TCGA-A2-A3XY | 786 | NA | Alive |
| TCGA-A7-A0DA | 373 | NA | Alive |
| TCGA-A7-A26G | 210 | NA | Alive |
| TCGA-A7-A4SE | 371 | NA | Alive |
| TCGA-A7-A6VV | 181 | NA | Alive |
| TCGA-A7-A6VW | 176 | NA | Alive |
| TCGA-A7-A6VY | 125 | NA | Alive |
| TCGA-A8-A07C | 580 | NA | Alive |
| TCGA-A8-A07O | 304 | NA | Alive |
| TCGA-A8-A07U | 303 | NA | Alive |
| TCGA-A8-A08R | 30 | NA | Alive |
| TCGA-A8-A09X | NA | 426 | Dead |
| TCGA-AC-A2BK | 1172 | NA | Alive |
| TCGA-AC-A2QJ | 69 | NA | Alive |
| TCGA-AC-A6IW | 21 | NA | Alive |
| TCGA-AC-A7VC | 1 | NA | Alive |
| TCGA-AN-A04D | 52 | NA | Alive |
| TCGA-AN-A0AL | 198 | NA | Alive |
| TCGA-AN-A0AR | 10 | NA | Alive |
| TCGA-AN-A0AT | 10 | NA | Alive |
| TCGA-AN-A0G0 | 16 | NA | Alive |
| TCGA-AN-A0XN | 10 | NA | Alive |
| TCGA-AN-A0XS | 10 | NA | Alive |
| TCGA-AN-A0XU | 10 | NA | Alive |
| TCGA-AO-A03U | NA | 1793 | Dead |
| TCGA-AO-A0J4 | 294 | NA | Alive |
| TCGA-AO-A0J6 | 775 | NA | Alive |
| TCGA-AO-A0JL | 1319 | NA | Alive |
| TCGA-AO-A124 | 3120 | NA | Alive |
| TCGA-AO-A128 | 2877 | NA | Alive |
| TCGA-AO-A129 | 2923 | NA | Alive |
| TCGA-AO-A12F | 1471 | NA | Alive |
| TCGA-AO-A1KR | 2141 | NA | Alive |
| TCGA-AQ-A04J | 160 | NA | Alive |
| TCGA-AR-A0TS | 1138 | NA | Alive |
| TCGA-AR-A0TU | 360 | NA | Alive |
| TCGA-AR-A0U4 | 1684 | NA | Alive |
| TCGA-AR-A1AR | NA | 524 | Dead |
| TCGA-AR-A1AY | 615 | NA | Alive |
| TCGA-AR-A256 | NA | 2854 | Dead |
| TCGA-AR-A2LR | 620 | NA | Alive |
| TCGA-AR-A5QQ | NA | 322 | Dead |
| TCGA-B6-A3ZX | 1132 | NA | Alive |
| TCGA-B6-A400 | 215 | NA | Alive |
| TCGA-B6-A402 | 1651 | NA | Alive |
| TCGA-BH-A0B3 | 1203 | NA | Alive |
| TCGA-BH-A0B9 | 1572 | NA | Alive |
| TCGA-BH-A0BG | 756 | NA | Alive |
| TCGA-BH-A0BL | 1340 | NA | Alive |
| TCGA-BH-A0E0 | 134 | NA | Alive |
| TCGA-BH-A0RX | 170 | NA | Alive |
| TCGA-BH-A0WA | 372 | NA | Alive |
| TCGA-BH-A18G | 61 | NA | Alive |
| TCGA-BH-A1EW | NA | 1694 | Dead |
| TCGA-BH-A1F6 | NA | 2965 | Dead |
| TCGA-BH-A1FC | NA | 3472 | Dead |
| TCGA-BH-A42U | 3324 | NA | Alive |
| TCGA-BH-A6R9 | 160 | NA | Alive |
| TCGA-C8-A12V | 0 | NA | Alive |
| TCGA-C8-A131 | 14 | NA | Alive |
| TCGA-C8-A1HJ | 5 | NA | Alive |
| TCGA-C8-A26X | 11 | NA | Alive |
| TCGA-C8-A26Y | 0 | NA | Alive |
| TCGA-C8-A27B | 30 | NA | Alive |
| TCGA-C8-A3M7 | 1 | NA | Alive |
| TCGA-D8-A13Z | 210 | NA | Alive |
| TCGA-D8-A143 | 172 | NA | Alive |
| TCGA-D8-A147 | 2 | NA | Alive |
| TCGA-D8-A1JF | 96 | NA | Alive |
| TCGA-D8-A1JK | 0 | NA | Alive |
| TCGA-D8-A1JL | 265 | NA | Alive |
| TCGA-D8-A1XK | 326 | NA | Alive |
| TCGA-D8-A1XQ | 177 | NA | Alive |
| TCGA-D8-A1XW | 118 | NA | Alive |
| TCGA-D8-A27F | 106 | NA | Alive |
| TCGA-D8-A27H | 19 | NA | Alive |
| TCGA-D8-A27M | 145 | NA | Alive |
| TCGA-E2-A14N | 1350 | NA | Alive |
| TCGA-E2-A14R | 845 | NA | Alive |
| TCGA-E2-A14X | 692 | NA | Alive |
| TCGA-E2-A150 | 591 | NA | Alive |
| TCGA-E2-A158 | 450 | NA | Alive |
| TCGA-E2-A1II | 850 | NA | Alive |
| TCGA-E2-A1L7 | 633 | NA | Alive |
| TCGA-E2-A1LH | 2876 | NA | Alive |
| TCGA-E2-A1LL | 1014 | NA | Alive |
| TCGA-E2-A1LS | 239 | NA | Alive |
| TCGA-E9-A5FL | 8 | NA | Alive |
| TCGA-EW-A1OV | 523 | NA | Alive |
| TCGA-EW-A1OW | 464 | NA | Alive |
| TCGA-EW-A1P4 | 501 | NA | Alive |
| TCGA-EW-A1P8 | NA | 239 | Dead |
| TCGA-EW-A1PB | 608 | NA | Alive |
| TCGA-EW-A1PH | 140 | NA | Alive |
| TCGA-EW-A3U0 | 229 | NA | Alive |
| TCGA-EW-A6SB | 760 | NA | Alive |
| TCGA-GI-A2C9 | 711 | NA | Alive |
| TCGA-GM-A2DB | 1616 | NA | Alive |
| TCGA-GM-A2DF | 1299 | NA | Alive |
| TCGA-GM-A2DH | 1286 | NA | Alive |
| TCGA-HN-A2NL | 79 | NA | Alive |
| TCGA-LL-A441 | 91 | NA | Alive |
| TCGA-LL-A5YO | 97 | NA | Alive |
| TCGA-LL-A73Y | 126 | NA | Alive |
| TCGA-OL-A6VO | 480 | NA | Alive |
| TCGA-S3-AA10 | 241 | NA | Alive |
